# *Wolbachia* uses ankyrin repeats to target specific fly proteins

**DOI:** 10.1101/2025.05.15.654400

**Authors:** William Hamilton, Jonathan Massey, Erin Hardy, Sergio López-Madrigal, Melissa Phelps, MaryAnn Martin, Irene Newton

## Abstract

Arthropods, the most diverse phylum on Earth, are hosts to a plethora of bacterial parasites that secrete various effectors of unknown function during infection. The most prevalent of these is the intracellular bacterium *Wolbachia pipientis*. The microbe infects between 40-60% of insect species, causes important reproductive manipulations, and limits virus replication in arthropod vectors, becoming a promising biocontrol agent. Understanding the molecular basis of *Wolbachia* infection and *Wolbachia-*induced phenotypes is critical to the use of *Wolbachia* in vector control. These *Wolbachia* ankyrin repeat proteins (WARPs) represent a highly dynamic and diverse part of the *Wolbachia* pangenome and remain thus far, largely uncharacterized. Here, we perform molecular and genetic screens to identify interactions between *Wolbachia w*Mel WARPs and their target host proteins in *Drosophila melanogaster*. Our results identify strong interactions of two *Wolbachia* proteins, WARP434 and WARP754, with two host targets (CG11327 and Ptp61F, respectively). Heterologous expression of these two WARPs is extremely toxic in *Drosophila* tissues and the toxicity is dependent on the ankyrin repeat domain of each WARP. Importantly, knockdown of the host targets alleviates toxicity, confirming WARP/target interactions. Finally, antibodies targeting both WARPs show expression by *Wolbachia* during infection of *Drosophila* cells. Understanding how *Wolbachia* manipulates its host biology and which host pathways it targets during infection will help us divine how the most prevalent intracellular bacterial parasite on Earth interacts with its insect hosts at the molecular level. Our screen is the first step towards that goal.

**Importance:** Molecular interactions drive co-evolutionary arms races between hosts and pathogens. These interactions shape the structure and function of both host and parasite proteins, enabling immunity or virulence during infection. Understanding the molecular details that unfold during these events illustrates not only how hosts and parasites co-evolve at the molecular level but also may help characterize the function of poorly understood proteins. The most prevalent intracellular infection on earth is Wolbachia pipientis, with between 40-60% of insects harboring the bacterial symbiont. Understanding how Wolbachia infects host cells and the molecular tools it uses to alter cell biology is critical to the use of the microbe in vector control. Here, we identify Wolbachia proteins used by the symbiont to interface with specific host proteins. Understanding the molecular mechanisms underlying this host-microbe interaction will shed light on how an important symbiont, used in the control of vector populations and disease transmission, uses WARPs to interact with host targets and how targeting this host protein contributes to infection.

## Introduction

*Wolbachia pipientis* is an obligate intracellular microbe and arguably the most successful infection on our planet, colonizing 40-60% of insect species (1, 2). *Wolbachia* are alpha-proteobacteria, part of the intracellular *Anaplasmataceae*, and related to the important human pathogens *Anaplasma, Rickettsia* and *Ehrlichia* (3). However, *Wolbachia* do not infect mammals, but instead are well known for their reproductive manipulations of insect populations, inducing phenotypes such as male-killing, feminization, or sperm-egg incompatibility (4). In the last decade, *Wolbachia* have also been shown to provide a benefit to insects, where the infection can inhibit RNA virus replication within the host (5), a phenomenon known as pathogen blocking. Because insects are vectors for disease, and *Wolbachia* alter the ability of these vectors to harbor important human pathogens, *Wolbachia* are being used to control the spread of arthropod-borne diseases such as dengue.

The establishment of the *Wolbachia* infection itself (i.e. host cell invasion, persistence, proliferation, and transmission to the next generation) is a prerequisite for the use of *Wolbachia* as a biocontrol agent. This leads us to ask: *how does Wolbachia manipulate host cell biology to establish an infection?* Like all intracellular bacteria, *Wolbachia* need to alter the host cell to invade and persist. Many microbes accomplish this via secretion systems, nanomachines that enable the microbes to directly transfer proteins from the bacterium into the cytosol of host cells. Virtually all *Wolbachia* strains sequenced to date encode a functional type IV secretion system (T4SS) (6-8), which is expressed by *Wolbachia* within its native host (7, 9). In *Wolbachia*, some predicted secreted effectors are co-regulated with the T4SS and for a handful of these proteins, heterologous T4SS assays support their secretion by the symbiont (10, 11). Identifying the specific proteins *secreted* by *Wolbachia* has been a primary goal of the field. These proteins, referred to as effectors, often act to manipulate or usurp host cell processes to promote bacterial infection (12, 13). These modes include (but are not limited to) attacking the host cell surface to form pores, inactivating host cytosol machinery to collapse the cytoskeleton, or entering the nucleus to manipulate host gene regulation (14). At each stage of attack, the bacterial effectors often interact directly and specifically with host proteins to perturb a biological process that enables pathogen entry into or defense from the host cell (15-21). Understanding how bacterial effectors function, therefore, has taught scientists not only how pathogens cause disease, but also how fundamental cell biological mechanisms work in healthy tissue (22). While effectors are encoded in bacteria, they act within eukaryotic cells and hence feature domains that share structural, functional, and sequence similarity with eukaryotic proteins (12, 13, 23, 24).

Ankyrin repeats (ANKs) are one of the most common domains found in eukaryotic proteins. They were first discovered as repeat sequences in yeast (*Saccharomyces cerevisiae*) proteins Swi6 and Cdc10 and the *Drosophila melanogaster* protein Notch (25). They are characterized by a repeating 33 amino acid motif (containing the conserved N-terminal residues G-X-TPLHLA) that folds into a helix–loop–helix–β– hairpin/loop structure (26). It was previously thought that ANKs are primarily restricted to eukaryotic proteins, but evidence suggests they are also prevalent among bacteria and virus genomes, especially in those microbes that interact with eukaryotes (27, 28). Ankyrin repeat domains often mediate protein-protein interactions and act as a scaffold for protein recognition (29). With the sequencing of the first *Wolbachia* genome from strain *w*Mel, which naturally infects *Drosophila melanogaster*, researchers noted the prevalence of ankryin repeat domains among encoded proteins (6). Subsequent sequencing of more strains from the *Wolbachia* genus revealed that these intracellular microbes dedicate upwards of 4% of their genome to ankyrin repeat containing proteins, a much higher proportion than in other bacterial genera (28, 30). Interestingly, the *Wolbachia* ankyrin repeat proteins across the *Wolbachia* genus are also highly dynamic. Thus, they are known to change size through expansion and contraction, or domain swapping and addition, and are likely horizontally transferred between strains with the aid of bacteriophages (31). Many in the field therefore hypothesize that *Wolbachia’s* ankyrin repeat proteins (WARPs) are involved in manipulating host biology, and are likely secreted via one of two secretion systems found in virtually all sequenced *Wolbachia* genomes (T1SS or T4SS); indeed, closely-related microbes use ankyrin repeat containing proteins as secreted effectors (27, 32-36). For example, the pathogen *Anaplasma phagocytophilum* secretes at least three ankyrin repeat containing proteins (*e*.*g*. AnkABC) through its T4SS (37), and AnkA is known to localize to the nucleus, modifying host gene expression (38-40).

Here, we present our study of the 25 WARPs found in strain *w*Mel, from *Drosophila* (Figure 1). We show first that these WARPs are likely secreted through bioinformatics predictive software. We then show that two WARPs cause severe phenotypes in the fly when overexpressed and that these phenotypes are dependent on the presence of the ankyrin domain. A yeast-2-hybrid screen of the *Drosophila* orfeome reveals the host targets to which these toxic WARPs bind and suppression assays in the fly confirm this interaction. We show that these WARPs are made by *Wolbachia* during an infection and immunohistochemistry suggests they are secreted outside of the bacterial cell. Understanding the molecular mechanisms underlying this host-microbe interaction will shed light on how an important symbiont, used in the control of vector populations and disease transmission, uses WARPs to interact with host targets and how targeting this host protein contributes to infection.

**Figure 1.**
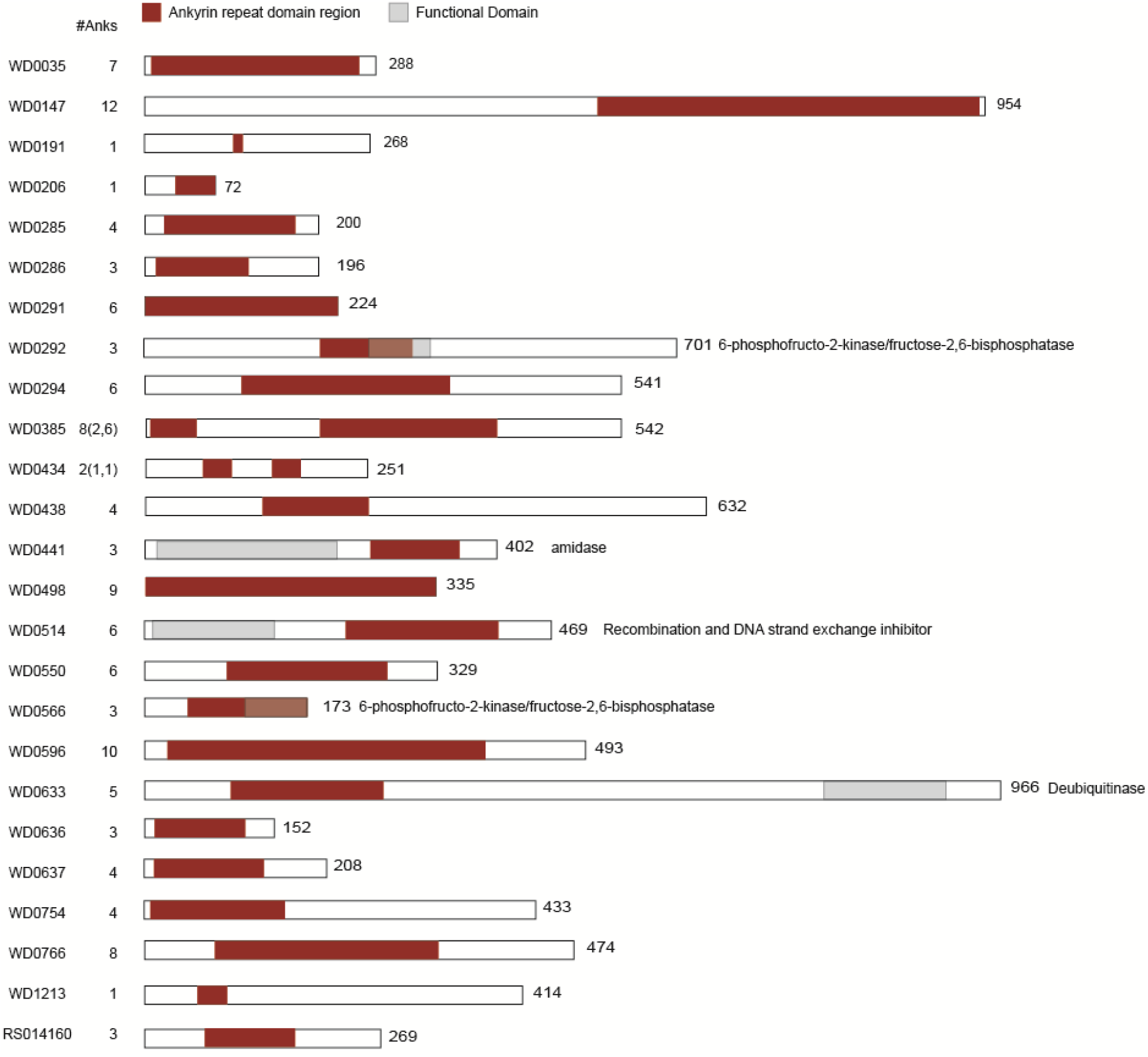
*Wolbachia* WARPs vary in length, number of ankyrin repeats, and functional domain content. For each *Wolbachia* WARP, the full protein length (right) and number of ankyrin repeats (left) identified by AnkPred software (41) are indicated. Ankyrin repeat boundaries are delineated in red. Predicted functional domains, shown in grey, were identified using HHPred searching the PFAM database (42).

## Results

### WARPs are predicted secreted effectors

The *Wolbachia* strain *w*Mel, a native symbiont of *Drosophila*, contains 25 WARP proteins (6), which vary in length from 72 to 966 amino acids and from 1 to 12 ankyrin repeats. (**Figure 1**). There is also significant variation in predicted functional domains associated with these WARPs with many of the proteins having no homology to any putative domain and others harboring wildly different putative enzymatic domains (such as amidases or glycolysis-associated enzymes). One of the WARPs, WD0633, encodes a deubiquitinase with homology to the OTU protein of *Drosophila melanogaster*.

For *Wolbachia* predicted effectors to act on host cell biology they must exit the bacterial cell via some mechanism. *Wolbachia* harbor both a T1SS and T4SS and therefore we sought to use bioinformatic prediction software to determine if any of the *w*Mel WARPs encode signatures of secretion. All 25 WARPs from the *w*Mel genome were used as input to the BastionX v.1.0 software suite (43)(fast mode, all functional analyses checked)(**Table 1**). A little more than half (16/25) of the WARPs were predicted to be secreted via the T4SS. By contrast, only 2/25 WARPs were predicted to be secreted by the T1SS. Few *Wolbachia* proteins have been tested for secretion using a heterologous assay in *E. coli* (11), and only a handful of secreted effectors have been visualized outside of the bacterial cell (WalE1, the *cif* proteins (44, 45)). This computational result, therefore, encouraged us to explore WARP-induced phenotypes in *Drosophila*.

**Table 1.** WARP Protein secretion predictions based on BastionX. Putatively secreted *Wolbachia* ankryin repeat containing proteins listed by accession and BastionX prediction results highlighted.

| Accession | Prediction Results based on Single Type Predictors |  |  |  |  | Prediction Results Total |  |
| --- | --- | --- | --- | --- | --- | --- | --- |
|  | Type I | Type II | Type III | Type IV | Type VI | Substrate | Possible Secretion System |
| >Query_WD_0035 | 0.215 | 0.195 | 0.475 | 0.663 | 0.275 | yes | IV |
| >Query_WD_0147 | 0.088 | 0.101 | 0.095 | 0.607 | 0.202 | yes | IV |
| >Query_WD_0191 | 0.105 | 0.138 | 0.154 | 0.405 | 0.053 | no | '- |
| >Query_WD_0206 | 0.054 | 0.147 | 0.337 | 0.328 | 0.126 | no | '- |
| >Query_WD_0285 | 0.057 | 0.042 | 0.284 | 0.455 | 0.109 | no | '- |
| >Query_WD_0286 | 0.029 | 0.038 | 0.096 | 0.345 | 0.099 | no | '- |
| >Query_WD_0291 | 0.542 | 0.289 | 0.62 | 0.655 | 0.613 | yes | I III IV VI |
| >Query_WD_0292 | 0.205 | 0.298 | 0.765 | 0.904 | 0.534 | yes | III IV VI |
| >Query_WD_0294 | 0.17 | 0.137 | 0.187 | 0.449 | 0.225 | no | '- |
| >Query_WD_0385 | 0.71 | 0.14 | 0.806 | 0.706 | 0.443 | yes | I III IV |
| >Query_WD_0434 | 0.084 | 0.217 | 0.396 | 0.557 | 0.085 | yes | IV |
| >Query_WD_0438 | 0.074 | 0.127 | 0.627 | 0.953 | 0.363 | yes | III IV |
| >Query_WD_0441 | 0.141 | 0.101 | 0.522 | 0.911 | 0.25 | yes | III IV |
| >Query_WD_0498 | 0.102 | 0.197 | 0.562 | 0.771 | 0.197 | yes | III IV |
| >Query_WD_0514 | 0.164 | 0.07 | 0.273 | 0.662 | 0.175 | yes | IV |
| >Query_WD_0550 | 0.179 | 0.113 | 0.126 | 0.231 | 0.145 | no | '- |
| >Query_WD_0566 | 0.042 | 0.112 | 0.219 | 0.554 | 0.108 | yes | IV |
| >Query_WD_0596 | 0.081 | 0.063 | 0.144 | 0.342 | 0.027 | no | '- |
| >Query_WD_0633 | 0.283 | 0.185 | 0.838 | 0.936 | 0.502 | yes | III IV VI |
| >Query_WD_0636 | 0.11 | 0.088 | 0.214 | 0.563 | 0.115 | yes | IV |
| >Query_WD_0637 | 0.08 | 0.112 | 0.392 | 0.681 | 0.144 | yes | IV |
| >Query_WD_0754 | 0.07 | 0.075 | 0.619 | 0.835 | 0.219 | yes | III IV |
| >Query_WD_0766 | 0.162 | 0.088 | 0.291 | 0.495 | 0.169 | no | '- |
| >Query_WD_1213 | 0.405 | 0.479 | 0.871 | 0.913 | 0.726 | yes | III IV VI |
| >Query_WD_RS04160 | 0.052 | 0.114 | 0.414 | 0.5 | 0.087 | no | '- |

### WARPs cause toxic phenotypes in Drosophila melanogaster

We reasoned that the WARPs, if interfacing with *Drosophila* biology, would induce phenotypes upon overexpression. We built or acquired UAS expression lines for all *w*Mel WARPs (see Methods). All WARP sequences were codon optimized for *Drosophila* and inserted into attP40 (chromosome II) or attP2 (chromosome III). Flies were homozygosed and crossed to GAL4 expression lines to determine toxicity upon induction. Global expression was compared to expression in the wing pouch and margin, the genitalia and sex combs, the dorsal midline, the eye, and the ovaries. Nearly all WARPs showed no evident phenotype at all upon expression (Figure S1), although we caution interpretation of this result, which could be related to timing and extent of protein expression. Importantly, expression of the well characterized *Wolbachia* effector WalE1 also shows no gross phenotype in *Drosophila* (8); lack of phenotype should not be interpreted as lack of function in the symbiosis. In our screen, two WARP constructs consistently produced highly toxic phenotypes upon expression in all tested drivers: WARP434 and WARP754 (Figure 2). Expression of WARP434 lead to death prior to eclosion when expressed globally, in the ovaries, or in the dorsal midline (only 4 female flies were able to eclose with the pictured incomplete thorax; no male flies eclosed). In contrast, expression in the eye led to a glassy-eye phenotype. When expressed in the wings (either margin or pouch) we observed blistering/crumpling of the adult wing that occurs from improper wing vein development. Finally, WARP434 expression caused ablation of both female and male genitalia and ablation of the male sex combs. Like WARP434, WARP754 also caused death when expressed globally, in the ovaries, or in the dorsal midline. WARP754 also induced blistering/crumpling wings when expressed in the wing pouch and ablation of the genitalia. However, unlike WARP434, WARP754 also caused death when expressed in the eye, and the phenotype observed upon expression in the wing margin was clipping on the distal edges of the wing. All the listed phenotypes but the wing pouch were highly prevalent, occurring in all or nearly all observed flies (Table 2).

**Table 2.** Ankyrin domains of WARPs mediate toxicity in flies upon expression. Counts of flies exhibiting toxic phenotypes (Figure 2) shown over total. In some cases no transgenic flies eclosed due to WARP-induced lethality (demarcated as DEAD).

|  | LongGMR-GAL4 | Vestigial-GAL4 | Apterous-GAL4 | Pannier-GAL4 | OvarianTumor-GAL4 | Doublesex-GAL4 | Act5C-GAL4 |
| --- | --- | --- | --- | --- | --- | --- | --- |
| **WARP 434** |  |  |  |  |  |  |  |
| male<br>(phenotype/total) | 102/102 | 107/109 | 37/173 | 0/DEAD | 0/DEAD | 108/116 | 0/DEAD |
| female<br>(phenotype/total) | 135/135 | 122/127 | 48/180 | 4/DEAD | 0/DEAD | 104/113 | 0/DEAD |
| **DAnk434** |  |  |  |  |  |  |  |
| male<br>(phenotype/total) | 0/109 | 0/115 | 0/102 | 0/107 | 0/144 | 0/110 | 0/109 |
| female<br>(phenotype/total) | 0/102 | 0/110 | 0/111 | 0/116 | 0/131 | 0/117 | 0/123 |
| **WARP 754** |  |  |  |  |  |  |  |
| male<br>(phenotype/total) | 0/DEAD | 132/132 | 17/102 | 0/DEAD | 0/DEAD | 104/107 | 0/DEAD |
| female<br>(phenotype/total) | 0/DEAD | 119/119 | 99/104 | 0/DEAD | 0/DEAD | 91/100 | 0/DEAD |
| **DAnk754** |  |  |  |  |  |  |  |
| male<br>(phenotype/total) | 0/100 | 0/103 | 0/106 | 0/101 | 0/112 | 0/107 | 0/148 |
| female<br>(phenotype/total) | 0/114 | 0/128 | 0/113 | 0/119 | 0/127 | 0/105 | 0/113 |

**Figure 2.**
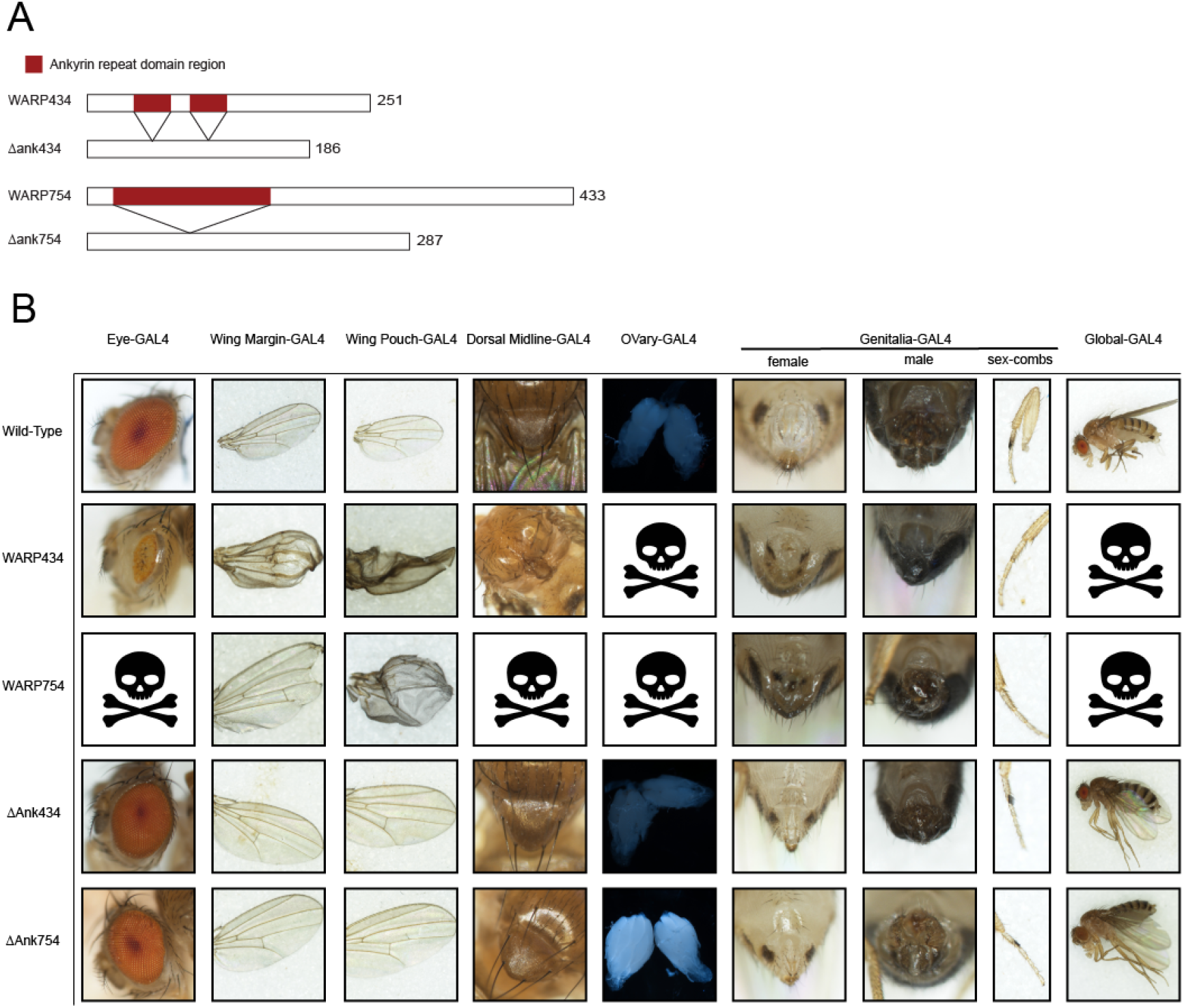
WARP434 and WARP754 induce toxic phenotypes in the fly upon expression. (A) Schematic of constructs generated for expression in the fly, both full length WARPs and those missing the ankyrin repeat domains. (B) Wild-type tissues (across top) compared to expression of constructs suggests that the ankyrin repeat domain containing regions of WARPs are required for toxicity.

Ankyrin repeat domains mediate protein-protein interactions and for bacterial effectors, are critical to bacterial targeting of specific host proteins (27, 46, 47). Without the ankyrin repeat domain, the bacterial effector no longer properly localizes (27, 47). Therefore, we hypothesized that the ankyrin domains of WARPs 434 and 754 might mediate the toxicity observed upon expression in *Drosophila*. We next generated constructs for both WARPs that lack the ankyrin repeat regions (Figure 2A). Interestingly, upon expression of the proteins lacking the ankyrin domains, we no longer observed toxicity, suggesting that the ankyrin repeat domain is necessary for targeting of specific host proteins (Figure 2B). Yet, it is possible that truncation of the protein abrogated proper folding. We therefore went on to directly test whether the ankyrin repeat domains of WARPs 434 and 754 directly bind *Drosophila* proteins.

### WARPs bind to specific host targets Ptp61F and CG11327

Our hypothesis was that the ankyrin repeat domains of WARPs mediate interactions with specific *Drosophila* proteins, thereby inducing the phenotypes observed in the whole animal upon expression. One way in which to test for direct binding is a yeast-2-hybrid assay. This assay relies on the reconstitution of a transcription factor in the yeast to drive expression of a metabolic marker gene, allowing for growth in hybrids that harbor interacting proteins. We generated truncated constructs to include primarily the ankyrin repeat domains of WARP434 and 754 (Figure 3A). These were fused to the DNA binding domain of GAL4 using the pDest-DB-cen plasmid and transformed into *S. cerevisiae* (Y8930). A pool containing both Y8930 DB-ankyrin strains for WARP434 and WARP754 was screened for autoactivation by mating to a pDest-AD-empty Y8800 strain. After 96 h at 30ºC, we found no evidence of growth on Y2H selective media containing 1mM 3AT (3-amino-1,2,4-triazole; Sigma-Aldrich A8056), indicating no autoactivation (48). These WARP-expressing yeasts were then mated to the *S. cerevisiae* activation domain (AD) library containing the Drosophila ORFeome, a collection of 10,490 AD-*Drosophila* ORF fusion strains representing ∼2/3 of the *D. melanogaster* proteome. Each stock was mated with the 10,168 viable Y8800 AD-*Drosophila* ORF strains (322 strains did not grow up from the glycerol stock; Table S1), plated on Y2H selective media containing 1mM 3AT, and grown at 30°C for 72-96 h.

**Figure 3.**
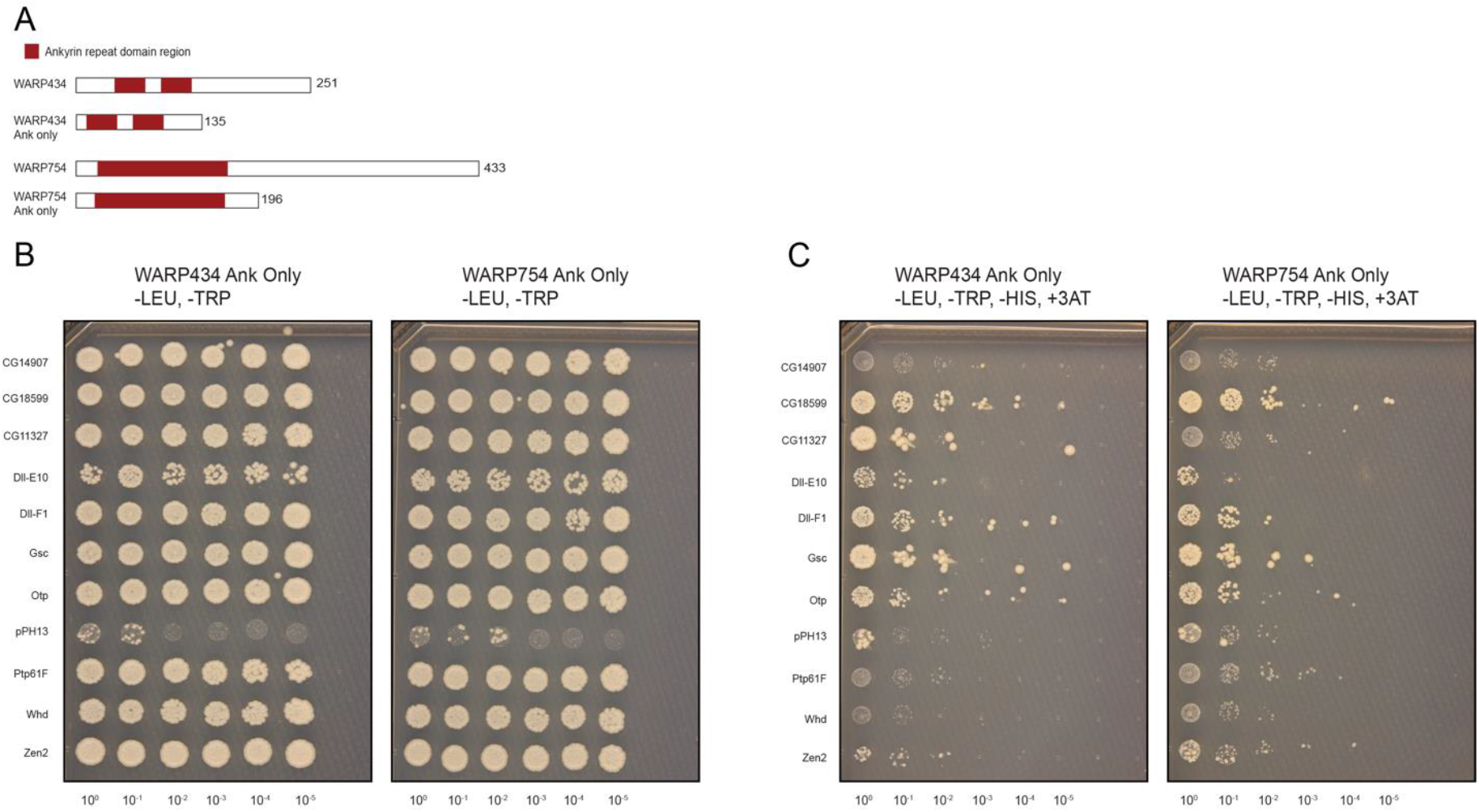
The ankyrin repeat domains of 434 and 754 interact with *Drosophila* proteins in a yeast-2-hybrid assay. (A) Schematic of constructs generated for expression in yeast. The N-terminal portions of the proteins containing the ankyrin repeat domains were included in the screen (B) Yeast hybrids grown under permissive conditions were (C) replica spotted onto media lacking histidine and containing the drug 3AT to screen for interactions. WARP434 interacts with CG11327 and WARP754 with Ptp61f.

The Y2H assay splits a transcription factor (typically GAL4 or LexA) into the DNA-binding domain (BD) and the activation domain (AD). If the WARP protein, fused to the BD interact with a *Drosophila* protein fused to the AD domain, the BD and AD are brought together, reconstituting a functional transcription factor. This drives expression of reporter gene (HIS3), allowing growth on selective media, in this case, media without histidine. We also used 3AT, a competitive inhibitor of the HIS3 gene product, imidazoleglycerol-phosphate dehydratase, which is involved in histidine biosynthesis. In the presence of 3AT, and when HIS3 is used as the reporter gene, yeast cells can grow on media lacking histidine only if HIS3 is sufficiently expressed. Using this methodology, we identified putative host target proteins for both WARPs 434 and 754 (Figure 3C). Several of our candidate interactors turned out to be auto-inducers (CG18599, DII, Gsc, Otp, pPH13, Zen2), leaving only two potential host targets: WARP434 interacts with the *Drosophila* protein CG11327 while WARP754 interacts with Ptp61F (Figure 3C).

The yeast-2-hybrid initial screen suggested interactions between our most toxic WARP proteins and *Drosophila* proteins. However, these interactions could be an artifact of the yeast expression system. We therefore sought to confirm the involvement of each host protein in the phenotypes induced by the WARPs using *Drosophila* genetics. We established stable lines expressing WARPs in the eye (434;LongGMR-GAL4) or wing (754;AP-GAL4) tissue. Constitutive expression of the WARPs in these tissues leads to severe fly phenotypes (Figure 2B). We reasoned that if CG11327 and Ptp61F were directly interacting with WARPs 434 and 754 (respectively), we should observe a genetic interaction upon knockdown of these loci using RNAi. We acquired flies from the TRiP project harboring a UAS construct driving expression of a short hairpin targeting either CG11327 or Ptp61F (see Table S2 for fly stocks). We then crossed these TRiP flies to our lines that constitutively express the WARPs in the eye or wing to induce knockdown of the host targets and resulting progeny were scored for severity of toxicity induced by accompanying WARP expression. We were able to track the correct genotypes because we used balancer chromosomes with dominant markers (Figure 4). Interestingly, we observed a partial recovery of the eye upon CG11327 knockdown (KD) in the WARP434 expressing line (Figure 4A) and an alleviation of the fly wing phenotype for WARP754 expressing flies upon knockdown of Ptp61F in the wing tissue (Figure 4B). For flies expressing WARP434 simultaneously with the CG11327 KD, we observed a consistent partial eye recovery across all progeny expressing both constructs. In comparison, the penetrance of toxicity observed upon WARP754 expression is significantly reduced in the presence of the Ptp61F knockdown; 56% of flies showed a severe or mild wing phenotype upon WARP754 expression and this number dropped to 16% and 20% in the RNAi KD. Indeed, the total numbers of wild-type flies emerging from these KD crosses was much higher than that observed for the WARP754 constitutive expression line (43% in the original line vs. 83% and 79% in the KD lines). These results strongly suggest that WARPs 434 and 754 do indeed interact with CG11327 and Ptp61F, respectively.

**Figure 4.**
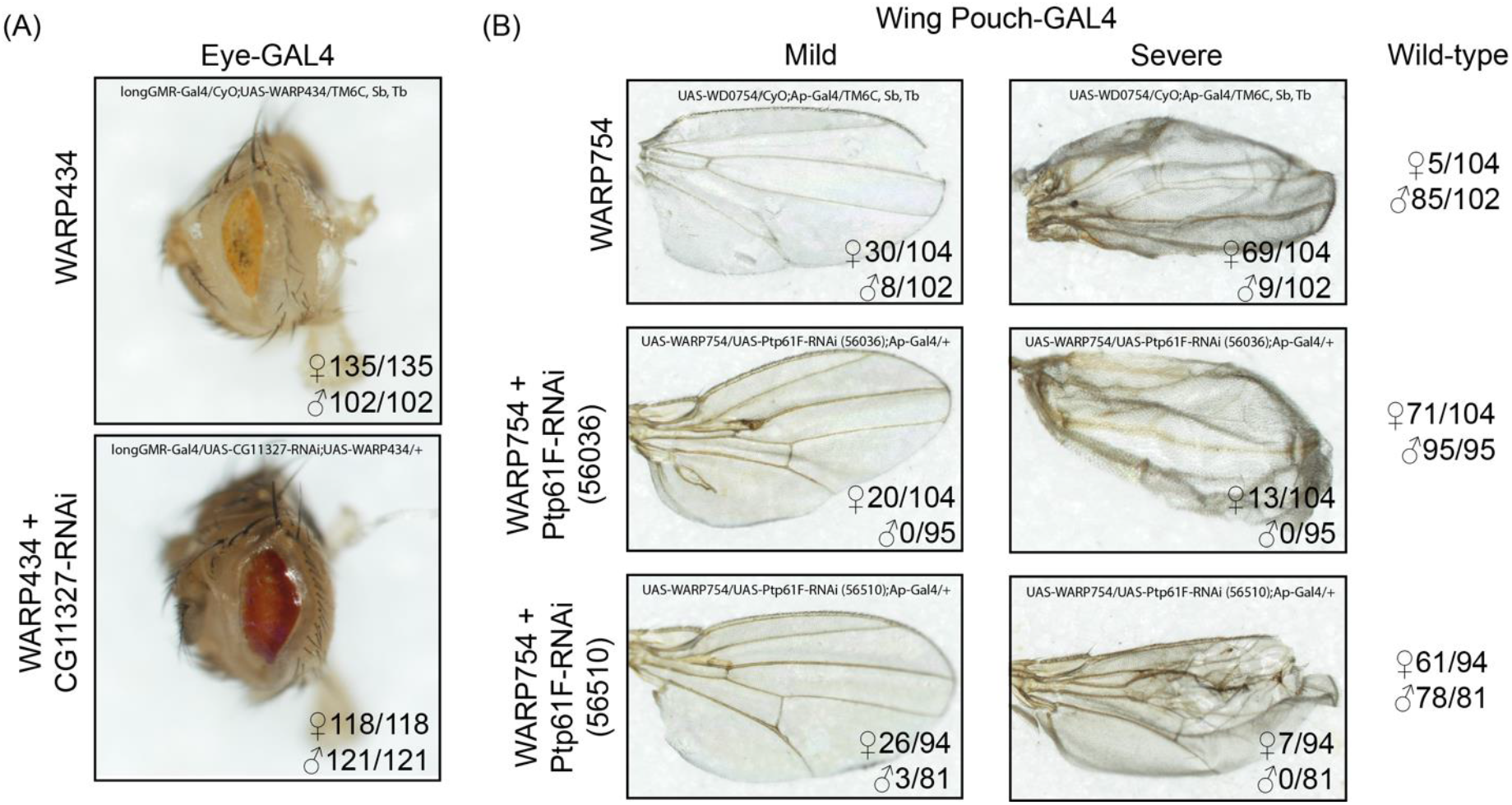
Concomitant knockdown of putative host targets for WARPs reduces observed toxicity in fly tissues. (A) Knockdown of CG11327 in flies expressing WARP434 recovers wild-type eyes. (B) Knockdown of Ptp61F with two distinct RNAi constructs in flies expressing WARP754 similarly recovers wild-type wings. Genotype of counted flies shown within each panel; flies were distinguished based on dominant phenotypic markers (*e*.*g*. Cyo, Tb, Sb).

### WARPs are expressed by Wolbachia in vivo

If WARPs 434 and 754 are used by *Wolbachia* during infection, we should be able to detect the protein in *Drosophila* cells infected by the bacterium. Towards that end, we generated polyclonal antibodies against full-length WARP434 and WARP754 and used the resulting sera to perform immunohistochemistry on a *Wolbachia*-infected *Drosophila* cell line and the uninfected counterpart (Figure 5). Both antibodies were non-reactive in *Wolbachia*-free cells (Figure 5, Mock) but highly reactive in the presence of *Wolbachia*. WARP754 seems to colocalize with DAPI staining of *Wolbachia* in infected cells but also to puncta separate from the microbe (Figure 5). WARP434 has a more diffuse staining pattern across the cell.

**Figure 5.**
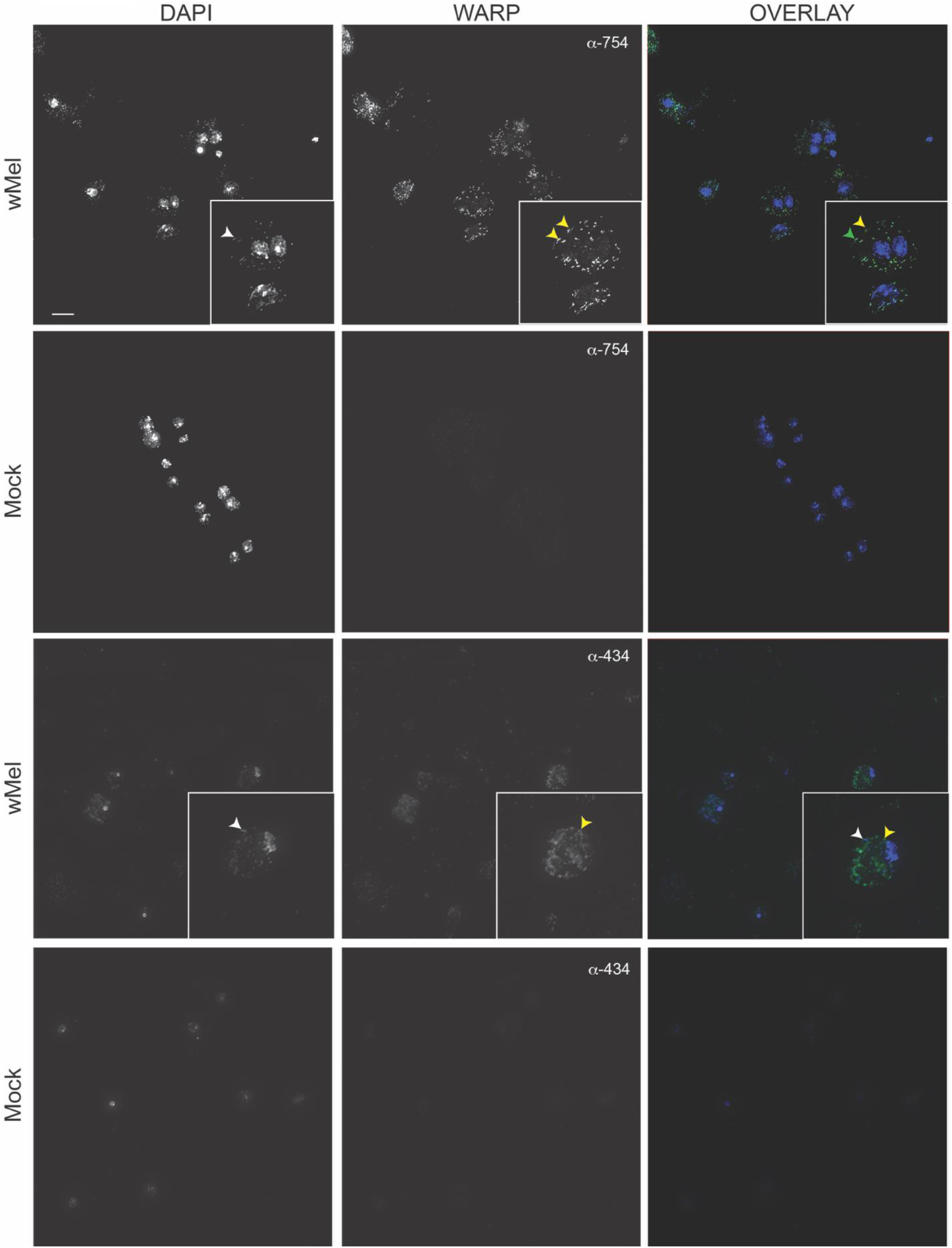
WARPs are expressed by *Wolbachia* during infection of *Drosophila* cells. Antisera against WARPs 434 and 754 detect expression by *Wolbachia* during infection of S2R+ cells. Only background staining is visible in uninfected cells (Mock). WARP754 staining (yellow arrowheads) seems to co-localize with *Wolbachia* DAPI staining (white arrowhead in DAPI panel; co-localization pointed to in green arrowhead in overlay figure) but also other foci within the cell. WARP434 staining is diffuse within the cell (yellow arrowhead) and does not seem to colocalize with *Wolbachia* DAPI puncta (white arrowhead). Scale bar 10 μm.

### WARP754 is toxic in adult flies

Expression of WARPs 434 and 754 during development is extremely toxic, limiting our ability to interrogate interactions between the expression of these proteins and the fly in the context of the *Wolbachia* symbiosis. We wondered if the toxicity was a result of expression at a specific developmental timepoint or if adult flies would similarly suffer from WARP toxicity. We therefore used the GAL80ts expression system to dampen expression from the GAL4 transcription factor, allowing recovery of adult flies, and induction of global expression after development. The GAL80ts works as a temperature-sensitive transcriptional repressor of GAL4; at cooler temperatures, it binds to GAL4 and prevents it from activating transcription. However, when temperatures are raised, GAL80ts loses its ability to bind GAL4, allowing for GAL4 to activate transcription. Flies containing GAL80ts and the UAS-WARP constructs were crossed to flies infected with *Wolbachia* and carrying the Actin-GAL4 driver. Crosses were kept at 18°C and progeny at the same temperature until eclosion. Adult flies were then sorted based on genetic markers and kept at 18°C, 23°C, or 30°C and survival of adults quantified. For both WARP constructs, we were able to acquire adult flies at 18°C. Flies expressing WARP434 are indistinguishable from controls when shifted to 23°C or 30°C, however, flies harboring the WARP754 construct quickly perished (Figure 6) and this phenotype was exacerbated by temperature. Additionally, for adult flies expressing WARP754 at 30°C, we observed a behavioral phenotype whereby adults lose the ability to control their movement (Supplemental Movies 1 and 2). This result suggests that WARP754 is toxic to flies, regardless of developmental stage expressed.

**Figure 6.**
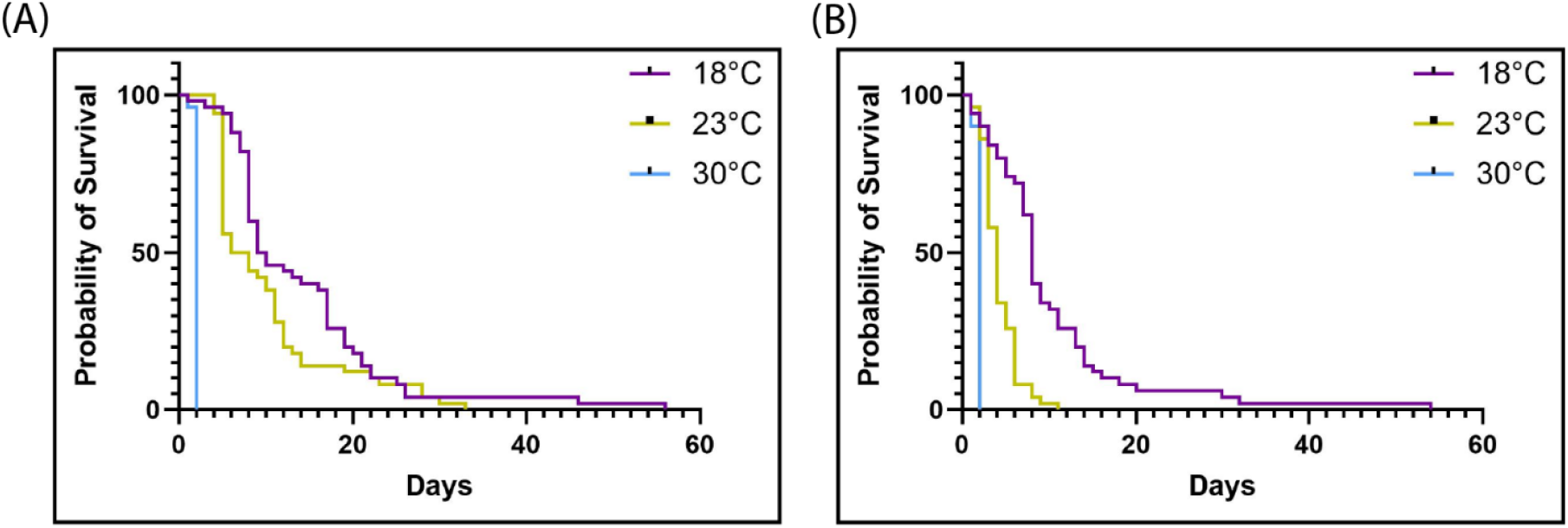
Adult flies quickly succumb when expression of WARP754 is induced. Flies were reared at 18°C under GAL80ts control of expression of the WARP constructs and shifted to 23°C or 30°C upon eclosure. For both (A) female and (B) male flies, expression of the construct resulted in a significant reduction in lifespan.

## Discussion

Here we present our discovery of novel host-microbe interactions in the *Wolbachia* symbiosis. The *Wolbachia* genus is a rich source of novel candidate effectors and bacterial toxins as the microbes are extraordinarily diverse and have proliferated across huge swathes of arthropod and nematode hosts. *Wolbachia* species harbor a huge number of ankyrin repeat domains (28), with some strains encoding upwards of 80 WARPs in a 1Mb genome. These proteins, therefore, will allow us to discover new aspects of cell biology in arthropods as the co-evolution of bacterial protein effectors and their targets has led to significant discoveries of basic cellular biology. For example, novel protein modifications such as AMPylation (49) were discovered by studying bacterial effectors. In the case of *Wolbachia*, the large number of WARPs harbored by this bacterial symbiont has captivated the imagination of scientists in the field (6), with many suggesting an involvement in reproductive parasitism (50-52). Indeed, WARPs are present in every *Wolbachia* genome sequenced to date and related microbes (*Anaplasma, Ehrlicia, Orientia, Rickettsia*) all use ankyrin repeat domain containing proteins to interact with their hosts and facilitate infection (53). Although the characterized effectors involved in cytoplasmic incompatibility (CifAB)(54), parthenogenesis (PifAB)(55) or male killing (wmk)(56) do not contain ankyrin repeat domains, that does not mean that all *Wolbachia* effectors are ankyrin free. Importantly, the male killing OSCAR gene recently identified from a *Wolbachia* strain infecting *Ostrinia scapulalis* moths is a WARP which targets the host Masculinizier protein to affect dosage compensation (57). Additionally, there is much about *Wolbachia* basic biology that has yet to be discovered, beyond the reproductive manipulations that have made the genus infamous; many secreted effectors are likely involved in surviving the host cell environment and facilitating transmission, as is the case for WalE1 (8, 44). Here, we show that two *Wolbachia*-encoded ankyrin repeat proteins (WARPs) cause severe toxic phenotypes upon overexpression and in combination with a yeast-2-hybrid screen, identify host targets of these WARPs. We use the toxic phenotype to confirm these targets through suppression of toxicity upon knockdown, suggesting that indeed, these are bona fide interactors of *Wolbachia* WARPs. This is the first large-scale characterization of *Wolbachia* ankyrin repeat domain containing proteins and their host targets.

What is known about these two *Wolbachia w*Mel-encoded WARPs? WARP434 is understudied in the field, with no publications we could find addressing its putative function or evolution in *Wolbachia*. In contrast, WARP754 has been studied in the context of expression in whole flies (58), as a putative secreted effector (10), for homology across *Wolbachia* and variation in ankyrin domain architecture, SNP content, and pseudogenization (31, 52, 59-61) and putative role in CI, for which none was found (50). Although WARP754 was previously reported to have homology to the IncA protein of *Chlamydia trachomatis* (7), no such homology could be supported using current databases. Although our staining suggests localization of WARP754 to the *Wolbachia*-containing vacuole, further work is required to interrogate WARP754’s localization and to determine if interaction with Ptp61F occurs on the vacuolar surface.

Our results suggest both that the ankyrin repeat domains of WARPs facilitate binding to *Drosophila* targets and that this binding is necessary for induced toxicity. We come to this conclusion because expression of the ankyrin domain alone recovered interactions with *Drosophila* proteins via yeast-2-hybrid and because the ankyrin deletion construct in flies did not induce any toxicity whatsoever (Figures 2,3). Many bacterial effectors use ankyrin repeats to bind to their host targets (46). For example, crystal structures of the *Legionella* effector AnkX bound to the Rab1 substrate suggest it uses its ankyrin repeats to mediate that specificity (62, 63) although importantly, associated domains in these multi-domain secreted effectors may also mediate toxicity (64). For WARPs 434 and 754, the ankyrin repeat domain seems to be sufficient for binding of CG11327 and Ptp61F (respectively) and necessary for induction of the toxic phenotype upon overexpression. Importantly, in our screen, we used the fly model to induce severe phenotypes and facilitate genetic dissection of WARP-host interactions; there is no evidence that these WARPs cause these specific phenotypes in these tissues when secreted by *Wolbachia*. Both CG11327 and Ptp61F expression is expected in tissues/organs colonized by *Wolbachia* (based on Flybase), but when and where these proteins are targeted by native WARPs 434 and 754 is still not known. In sum, the evidence presented here from WARP bioinformatic predictions, microscopy evidence for expression, and protein-protein interaction, suggest that these two host proteins are relevant for *Wolbachia*-host symbiosis.

Like WARP434 and WARP754, the targets of these two ankyrin repeat proteins are similarly understudied. CG11327, the target for WARP434, is an uncharacterized *Drosophila* gene for which few publications exist. It has been described as having weak similarity to transcription factors (65) but beyond this, little else is known regarding its function. For Ptp61F, because of its homology to mammalian proteins, we know a bit more about its putative functions. It is characterized as a non-receptor protein tyrosine phosphatase that negatively regulates JAK-STAT, insulin-like receptor, EGFR, and Pvr pathways impacting fly fecundity, growth, and lifespan (66), organ size (67), but also coordination of the actin cytoskeleton (68). These functional connections are particularly exciting and relevant for *Wolbachia* as infection by the microbe has been associated with changes in *Drosophila* fecundity and lifespan, as well as metabolism and the actin cytoskeleton (69-71) and our results provide a clear target for future mechanistic work determining *how* the microbe alters cell biology to achieve this organism-wide effect. Does WARP754 modify Ptp61F or alter its localization? Does WARP754 inactivate the protein or increase its activity? And what are the ramifications for *Wolbachia?* These questions are the subject of future work in our laboratory.

## Conclusion

We’ve shown that *Wolbachia* WARPs induce fly phenotypes and bind directly to fly proteins. This work identifies direct interactions between *Wolbachia* ankyrin repeat proteins and host targets, setting the stage for more mechanistic dissection of this important symbiosis.

## Methods

### Fly stocks, generation of UAS lines, and maintenance

All flies were grown at 23ºC with a 12 h light-dark cycle with lights on at 9AM and off at 9PM on standard corn-meal fly medium. For a full list of files used in this study see Table S2. Each open reading frame (ORF) predicted to encode at least one ankyrin repeat motif (HHpred and AlphaFold citation) from the *Wolbachia w*Mel strain genome (REF) (WD_0035, WD_0147, WD_0191, WD_0206, WD_0285, WD_0286, WD_0291, WD_0292, WD_0294, WD_0385, WD_0434, WD_0438, WD_0441, WD_0498, WD_0514, WD_0550, WD_0566, WD_0596, WD_0633, WD_0636, WD_0637, WD_0754, WD_0766, WD_1213, RS04160) was codon optimized for expression in *Drosophila melanogaster* in Geneious V2021.2 and synthesized at Genewiz Inc. with 5’ and 3’ Gibson tails compatible with plasmid pJFRC7 Table S2, with the exception of WD_0633, which was sent as a gift from Dr. Seth Bordenstein (Vanderbilt University) in a *D. melanogaster* expression plasmid containing a *D. melanogaster* codon optimized version of ORF WD_0633. The 20X UAS expression vector pJFRC7 (Addgene Plasmid #26220) was Midi prepped (Qiagen Cat. No. / ID: 12843) and digested with XhoI and XbaI to remove the mCD8::GFP insert. The resulting 8,145 bp band was separated on a 1% agarose gel and gel extracted (NEB Cat. No. T1020S) for Gibson Assembly (ThermoFisher Cat. No. A46624). Due to cloning difficulties from highly repetitive sequences, some *w*Mel ankyrin repeat genes (WD_0147, WD_0438, WD_0596, WD_0636, WD_1213) were cloned into the 20X UAS expression vector pPMW-attB (Addgene Plasmid #61814) via Gateway cloning. Briefly, synthesized dsDNA gene fragments were designed with 5’-CACC to facilitate ligation with the pENTR vector (ThermoFisher Cat. No. K240020). Products were transformed into chemically competent OneShot Top10 *E. coli* (ThermoFisher Cat. No. C404003), and three positive colonies were Mini prepped (Qiagen Cat. No. / ID: 27106) and Sanger sequenced to confirm the presence of each insert. Positive pENTR plasmids were combined with LR Clonase II Enzyme Mix (ThermoFisher Cat. No. 11791020) and the destination vector pPMW-attB, transformed and screened as described above. All positive plasmids were shipped to Rainbow Transgenics Inc. for injection into *D. melanogaster* embryos expressing the integrase PhiC31 to facilitate attP-attB integration into landing site attP40 (chr II) or attP2 (chr III). After injected P0 flies eclosed, they were crossed to siblings, and red-eyed F1 progeny were selected and crossed to keep as homozygous lines. All plasmid and Genewiz synthesized gene sequences can be found in Supplementary File 1.

### Ankyrin deletion lines

Ankyrin deletion 20X UAS expression constructs for ORFs WD_0434 and WD_0754 were created to test the necessity of ankyrin repeats in host toxicity. To identify the N- and C-terminal bounds of reach ankyrin repeat, we aligned the conserved 33 amino acid ankyrin repeat motif NGRTPLHLAARNGHLEVVKLLLEAGADVNAKDK (72) with WD_0434 and WD_0754 amino acid sequences. To confirm the presence of each repeat motif, protein structure predictions were generated via AlphaFold (73, 74), and the resulting .PDB files were submitted to AnkPred (41). We identified two ankyrin repeat motifs in WD_0434 and four ankyrin repeat motifs in WD_0754. DNA sequences with these motifs removed (Supplementary File 1) were codon optimized for expression in *D. melanogaster* in Geneious V2021.2, synthesized, cloned into pJFRC7, and injected into *D. melanogaster* as described above.

### Yeast two hybrid

Yeast-two hybrid (Y2H) assays were carried out using strains, media, and techniques described in (48). Briefly, due to toxicity, ORFs containing just the ankyrin repeat motifs from WD_0434 and WD_0754 were Gateway cloned into the destination vector pDest-DB-cen to create N-terminal fusion constructs of the *Saccharomyces cerevisiae* GAL4 DNA binding domain (DB) with C-terminal ankyrin repeats (Supplementary File 1). Primers (Integrated DNA Technologies) containing 5’-CACC to facilitate ligation with the pENTR vector (ThermoFisher Cat. No. K240020) were designed to amplify the ankyrin repeat regions from our *D. melanogaster* codon optimized 20X UAS pJFRC7 constructs (described above). The resulting amplicons were column purified (Qiagen Cat. No. / ID:28104) and cloned into the pENTR vector (ThermoFisher Cat. No. K240020). Products were transformed into chemically competent OneShot Top10 *E. coli* (ThermoFisher Cat. No. C404003), and three positive colonies were Mini prepped (Qiagen Cat. No. / ID: 27106) and Sanger sequenced to confirm the presence of each insert. Positive pENTR plasmids were then combined with LR Clonase II Enzyme Mix (ThermoFisher Cat. No. 11791020) and the destination vector pDest-DB-cen and again screened via Sanger sequencing for each insert. The resulting pDest-DB-cen ankyrin constructs were then transformed into the yeast *S. cerevisiae* strain Y8930 and screened using techniques described in (48).

The initial Y2H primary screen was designed to identify *D. melanogaster* protein-protein binding partners for the ankyrin repeat motifs in WD_0434 and WD_0754. Dr. David Hill and Kerstin Spirohn (Dana-Farber Cancer Institute) kindly gifted the *S. cerevisiae* activation domain (AD) library (*Drosophila* ORFeome) as a collection of 118 glycerol plates containing 10,490 AD-*Drosophila* ORF fusion strains (Supplementary File 1) representing ∼2/3 of the *D. melanogaster* proteome (https://flybi.hms.harvard.edu/). A pool containing both Y8930 DB-ankyrin strains for WD_0434 and WD_0754 was screened for autoactivation by mating to a pDest-AD-empty Y8800 strain. After 96 h at 30ºC, we found no evidence of growth on Y2H selective media containing 1mM 3AT (3-amino-1,2,4-triazole; Sigma-Aldrich A8056), indicating no autoactivation (48). Using an Integra Biosciences Assist Plus pipetting robot, the pool was then mated against 10,168 Y8800 AD-*Drosophila* ORF strains (322 strains did not grow up from the glycerol stock; Table S1) plated on Y2H selective media containing 1mM 3AT, and grown at 30°C for 72-96 h. AD strains that showed signs of growth when mated with the ankyrin DB pool were then re-mated in a secondary screen against the separated Y8930 DB-ankyrin strains to identify putative binding partners for WD_0434 and WD_0754 ankyrin repeat motifs.

### Drosophila melanogaster cell culture

*Drosophila melanogaster* embryonic S2R+ cells (DGRC Stock 150; https://dgrc.bio.indiana.edu//stock/150; RRID:CVCL_Z831) were incubated with *Wolbachia pipientis* strain *wMel* isolated from infected JW18 cultured cells (75) or underwent a mock infection with host cellular material from tetracycline-treated, uninfected JW18-TET cells (44). JW18 or JW18-TET cells were pelleted (1000 g for 5 minutes) and resuspended in cell culture media to a density of 1.6 x 10^7^ cells/mL for each. Four mL of each cell type were subjected to bead lysis; 100 µL of 0.5 mm sterile glass beads (#11079105, BioSpec Products) vortexed on full three times, for three seconds each pulse. The bead-beaten cells were strained through sterile 70 micron filters (Corning: Falcon, #352350) and loaded onto five micron filter units (Millipore #UFC40SV25, Ultrafree®-CL Centrifugal Filter, 2 mL volume) for centrifugation at 5000 g for 10 minutes. The supernatants were discarded, and pellets were resuspended in one mL of culture medium. To infect or mock-infect S2R+ cells, five mL of S2R+ cultures of 3 x 10^6^ cells/mL were incubated for one week with either one mL of lysed JW18 resuspended pellet or 1 mL of lysed JW18-TET resuspended pellet, respectively. After one week, the cells were diluted 1:5 in fresh media and allowed to grow another week. The infection status was confirmed after a week by immunochemistry using a rabbit polyclonal antibody to full-length FtsZ after one passage by methods described in Martin, *et al*., 2024 (44). Cells with and without *Wolbachia* were maintained in T25 unvented cell culture flasks, in a dark drawer at room temperature in Schneider’s insect medium with L-glutamine and sodium bicarbonate, supplemented with 10% heat-treated fetal bovine serum and 1% penicillin/streptomycin solution.

### Antibody production

Full-length *w*Mel WD0754 (WARP754) and WD0434 (WARP434) protein-encoding constructs were synthesized by GenScript using codons optimized for *E. coli* and subcloned into pET30a vector with a 6xHis tag at the N terminus, in BL21 Star (DE3). Cultures were grown in LB medium containing kanamycin at 37C and 200 rpm. Once cell densities reached OD600 = 0.6-0.8, 0.5 mM IPTG was introduced for induction and cells were shifted to 15C for 16 hours. The proteins were purified using Ni-NTA columns (Sigma-Aldrich) and subsequently sent to Cocalico Biologicals for antibody generation. Pre-inoculation sera from 4 rabbits was screened on blots of whole fly protein with and without infection and the rabbits with lowest background reactivity chosen for inoculation. Rabbit polyclonal antibody sera against both full-length purified proteins were generated (Cocalico Biologicals, Inc) and used separately for immunohistochemistry (see below).

### Drosophila melanogaster cultured cell immunochemistry and microscopy

The *Drosophila melanogaster* S2R+ cells with *Wolbachia* strain *w*Mel, were used to visualize the location of antibodies to native WARP434 and WARP754. Mock-infected S2R+ cells were passed with 3.3 μg/μl tetracycline added to the culture medium and were used as an uninfected control cell line. Monolayers grown to confluence were harvested from 25 cm^2^ non-vented tissue culture flasks, and cells were counted using disposable hemacytometers (Fisher Scientific). Cells were overlaid as 100 μl of 2 x 10^6^ or 200 μl of 1 x 10^6^ cells onto Concanavalin-A coated 22 mm square No. 1.5 coverslips (ConA 0.5 mg/ml applied and dried on to sterile, acid-washed coverslips and leaving a 2 mm ConA-free border) (76, 77). Cells were allowed to settle, attach, and grow for two nights. Cell-coated coverslips were transferred to 6-well tissue culture plates, one coverslip/well, cell-side facing up, and the coverslips were washed twice in 500 μl of 1X PBS before cells were fixed in 4% paraformaldehyde in 500 μl PBS for 20 minutes, followed by four washes of 500 μl 1X PBST (0.2% Tween-20 added to 1X PBS).

Coverslips were then incubated in 1 mL blocking solution (PBST 0.2% Tween-20 and 0.5% BSA) for one hour at room temperature before replacing the block solution with primary antibody dilutions overnight in blocking solution at 4°C . Primary antibodies to *w*Mel proteins were rabbit anti-WD0434 preadsorbed against PVDF-blotted uninfected S2R+ total protein lysate and used at 1:500 dilution; and rabbit anti-WD0754 purified with Thermo Scientific™ Melon™ Gel IgG Spin Purification Kit # 45206 and used at 1:500 dilution. In the morning, coverslips were washed three times for five minutes each with one mL of 1X PBST and then allowed to incubate for two hours at room temperature in the dark in 100 μl of secondary antibodies diluted 1:1000 in the blocking solution (Thermo-Fisher A-31573 Donkey anti-Rabbit IgG (H+L) conjugated to Alexa Fluor™ 647). Coverslips were washed for five minutes three times with one mL PBST, followed by a final dip in a 500 mL beaker of distilled water. Excess moisture was wicked off the edge of the coverslip with a tissue followed by mounting in 10 μl of Prolong Gold Antifade Reagent with DAPI (Invitrogen™, P36935) per coverslip on glass slides. Images were taken as Z-series stacks at 0.2 micron intervals using a Nikon Ti2 fluorescent microscope with 100x oil objective and processed using NIS Elements software (Nikon). Exposure times and stack intervals were equivalent across compared experimental conditions.

## Acknowledgements

We thank Dr. Sarah Boothman for providing the *Wolbachia w*Mel infected S2R+ cell line. The S2R+ cell line was obtained from the Drosophila Genomics Resource Center (NIH Grant 2P40OD010949). Stocks obtained from the Bloomington Drosophila Stock Center (NIH R40OD018537) were used in this study. This work was supported by an NIH NIAID R01 award to ILGN (R01AI178815).

